# Osmotic pressure induces unexpected relaxation of contractile 3D microtissue

**DOI:** 10.1101/2025.02.25.640067

**Authors:** Giovanni Cappello, Fanny Wodrascka, Genesis Marquez-Vivas, Amr Eid Radwan, Parvathy Anoop, Pietro Mascheroni, Jonathan Fouchard, Ben Fabry, Davide Ambrosi, Pierre Recho, Simon de Beco, Martial Balland, Thomas Boudou

**Author notes:** Correspondence should be addressed to G.C. and T.B.

## Abstract

Cell contraction and proliferation, matrix secretion and external mechanical forces induce compression during embryogenesis and tumor growth, which in turn regulate cell proliferation, metabolism or differentiation. How compression affects tissue contractility, a hallmark of tissue function, is however unknown. Here we apply osmotic compression to microtissues of either mouse colon adenocarcinoma CT26 cells, mouse NIH 3T3 fibroblasts, or human primary colon cancer-associated fibroblasts. Microtissues are anchored to flexible pillars that serve as force transducers. We observe that low-amplitude osmotic compression induces a rapid relaxation of tissue contractility, primed by the deformation of the extracellular matrix. Furthermore, we show that this compression-induced relaxation is independent of the cell type, proportional to the initial tissue contractility, and depends on RhoA-mediated myosin activity. Together, our results demonstrate that compressive stress can relax active tissue force, and points to a potential role of this feedback mechanism during morphogenetic events such as onco- or embryogenesis.

## Introduction

Tissue growth and function are driven by complex interactions between its constituent cells, extracellular matrix (ECM), and external stimuli [1]. Besides biochemical signals, cells receive and respond also to physical cues such as mechanical forces, fluid pressure, or the stiffness of the extracellular matrix [2, 3]. Cells respond to these external stimuli e.g. by modulating their contractility, proliferation, metabolism or differentiation [1, 4, 5].

These mechano-biological feedback mechanisms are important for cancer progression [6, 7]. For example, forces generated by cancer-associated fibroblasts (CAF) remodel the tumor stroma and can promote tumor cell invasiveness [8, 9], whereas the amount and stiffness of the extracellular matrix that often surrounds solid tumors facilitates the buildup of solid-stress pressure generated by the proliferation of cancer cells and can limit or inhibit tumor growth [10].

In vitro investigations of patho-physiological growth under pressure ranging from 0.5 to 20 kPa, using confinement [11, 12] or osmotic pressure [13, 14], have demonstrated the crucial role of both the cytoskeletal machinery and the surrounding ECM in the response to compression or ECM reorganization [10, 15–18]. For instance, the regulation of proliferation by external pressure has been shown to depend on the tissue microstructure and mechanics, as growth is inhibited when compression pressure applies to both cells and ECM, whereas it remains unaffected when compression applies only to cells [16, 19].

As cells assess the mechanics of their environment by applying forces to it, the regulation of cell contractility by either proliferation-induced or CAF-generated solid stress may be key to understanding the feedback mechanism between solid stress and tumor growth. Indeed, the mechanical state of cells drives the ECM remodeling and stiffening [17, 18, 20], as well as cell proliferation [21–24], which in turn increases internal solid stress when it occurs in a confined environment. However the difficulty of assessing cell forces in 3D environments under compression leaves a notable gap in our understanding of its consequences on cell contractility.

While the active regulation of stress in microtissues under tension has been largely investigated [17, 25–28], in this work we focus on the regulation of contractile active stress in tissues under compression. We engineer 3D microtissues, i.e. microscale constructs of cells embedded in extracellular 3D collagen matrix, where the microstructure of the exogenous collagen is remodeled in fibers by the cells themselves [29, 30]. The living material is suspended between two flexible cantilevers, whose deflection gives direct access to tissue contractility in real time [31–33]. The microtissues are then osmotically compressed using a culture medium supplemented with dextran, which is biologically inert and therefore exerts a persistent osmotic stress equivalent to an external mechanical compressive stress of identical magnitude [34].

To delineate the respective roles of cells and collagen in the mechanical response to pressure, we applied selective compression to both cells and extracellular matrix, or to individual components by modulating the molecular weight of dextran in the culture medium [19, 35]. Although an osmotic pressure leads to a compression stress, due to dehydration, we observe a counter-intuitive elongation of 3D tissues upon pressure. This compression-induced elongation only happens in 3D tissues composed of ECM and actively contracting cells, and only upon global compression, i.e. when both components are mechanically loaded. Furthermore, we demonstrate that this mechanical response of tissues to compression is cell type-independent but proportional to the initial tissue contractility. Together, these results highlight a unique approach to examine the effects of compression on 3D tissues and evidence the complex mechanical feedback loop between external pressure, ECM deformation and cell contractility.

## Results

### Regulation of tissue contractility upon compression

To engineer microtissues, we use arrays of 800×400×200 µm wells within a PDMS mold. We pipette a suspension of cells (mouse colon adenocarcinoma CT26 cells, NIH3T3 fibroblasts or human primary cancer-associated fibroblasts) and monomeric neutralized collagen I in these microwells (Fig. 1a). Once polymerized, the collagen is compacted by the cells. Two T-shaped, flexible cantilevers incorporated within each microwell anchor the contracting collagen matrix, leading to the formation of a microtissue suspended between the top of the pair of cantilevers. The forces generated by the cells and acting on the surrounding collagen matrix result in a global tissue force that deflects the flexible cantilevers (Fig. 1a). Using linear bending theory and experimental measurements, we calibrated the spring constant of the cantilevers, which was then used to relate the measured cantilever deflections to the associated tissue-generated force [31–33].

**Fig. 1.**
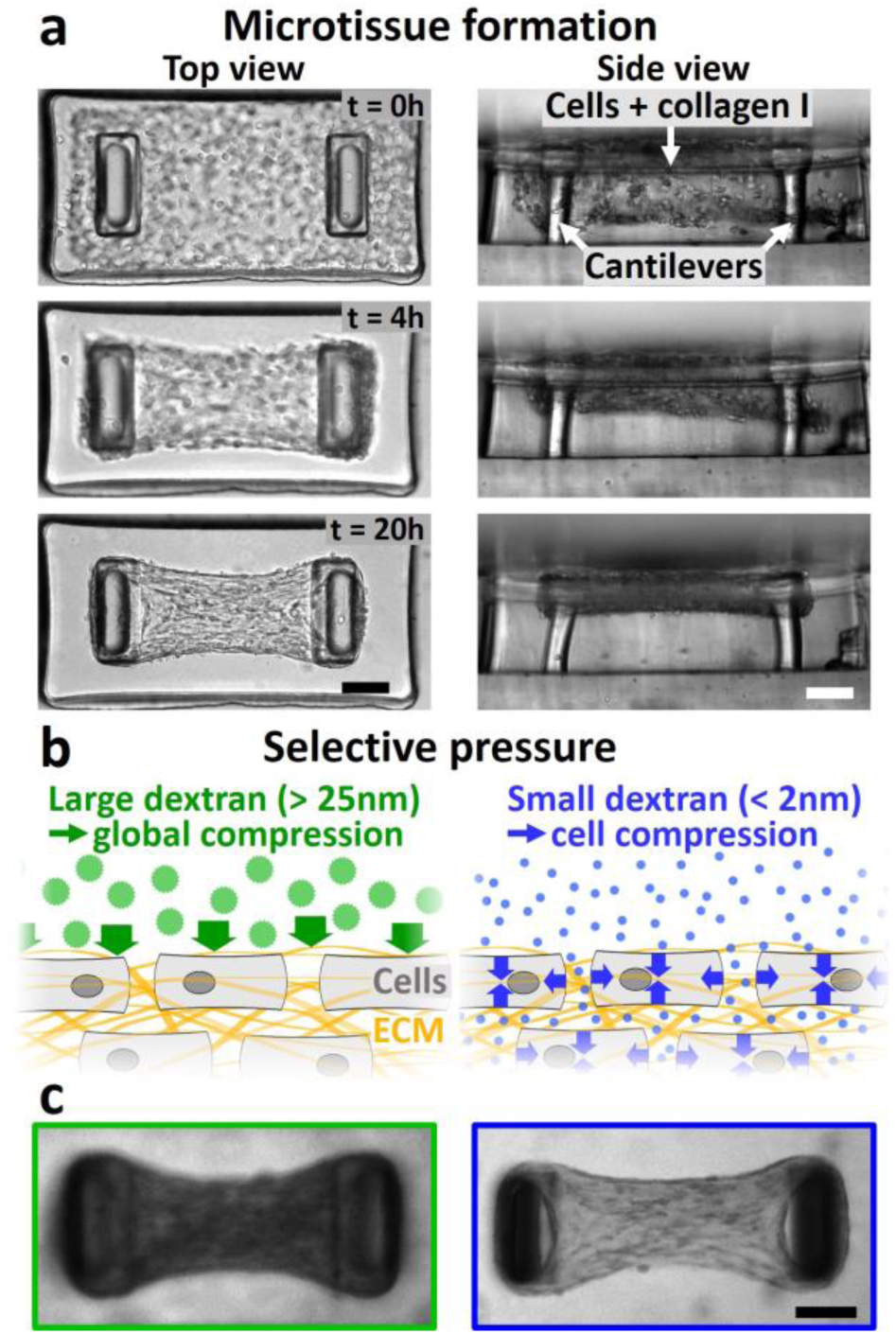
Microtissue engineering and selective compression. (a) Representative images of top view (left column) and side view (right column) illustrating the formation of a microtissue composed of NIH3T3 fibroblasts in collagen. Over time, the collagen matrix is spontaneously compacted by the fibroblasts to form a 3D microtissue suspended between T-shaped cantilevers. The deflection of the cantilevers, clearly visible after 20 h of formation, allows direct access to the tissue contractility. (b) Schematic of cells (gray) embedded in collagen (yellow fibers) submitted to selective compression. Large dextran osmolytes (green discs) do not permeate collagen and induce a global tissue compression (represented by green arrows), whereas small dextran osmolytes (blue dots) infiltrate collagen and induce a cell compression only (represented by blue arrows). (c) Fluorescently labeled large dextran molecules (gyration radius > 15 nm, left image) are excluded from the microtissue whereas fluorescently labeled small dextran molecules (gyration radius < 2 nm, right image) permeate the microtissue. Scale bars are 100 µm.

Once the tissue is formed and the produced force is stabilized, we used dextran osmolytes to compress the microtissue. Dextran macromolecules are biologically inert polysaccharides that do not readily enter the cytoplasm. Depending on their molecular weight - and therefore their size - they may be unable or only partially able to diffuse into the interstitial space. Consequently, smaller dextran molecules remain mostly in the extracellular space, resulting in an extracellular-intracellular concentration gradient, whereas larger dextran molecules stay predominantly outside the tissue, resulting in an extra tissue-interstitial gradient [13, 19, 35, 36]. In both cases, these gradients induce a persistent osmotic stress [13, 37]. Previous studies have established that 2 MDa dextran molecules with a gyration radius larger than 25 nm [38] do not permeate the cell-compacted collagen and thus compress the whole tissue, whereas small, 10 kDa dextran molecules with a gyration radius below 2 nm [38] infiltrate the collagen network and compress the cells only (Fig. 1b-c) [19, 35].

When applying a tissue compression Π = 1 kPa to NIH 3T3 microtissues using large dextran, we observed a transient, transverse (i.e. perpendicular to the long axis of the tissue) compression followed by a tissue elongation of 3.5 ± 1.1 % (12.7 ± 3.8 µm ) (Fig. 2a and Supp. Movie 1). The force as measured by the pillar deflection decreased from its initial baseline level F_0_ = 8.4 ± 1.9 µN to F_Π_ = 5.5 ± 1.6 µN in approximately 15 min (Fig. 2b). This tissue elongation and force relaxation was reversible, as tissue force increased back to its initial level after dextran removal, but at a slower rate, in approximately 60 min (Supp. Fig. 1).

**Fig. 2.**
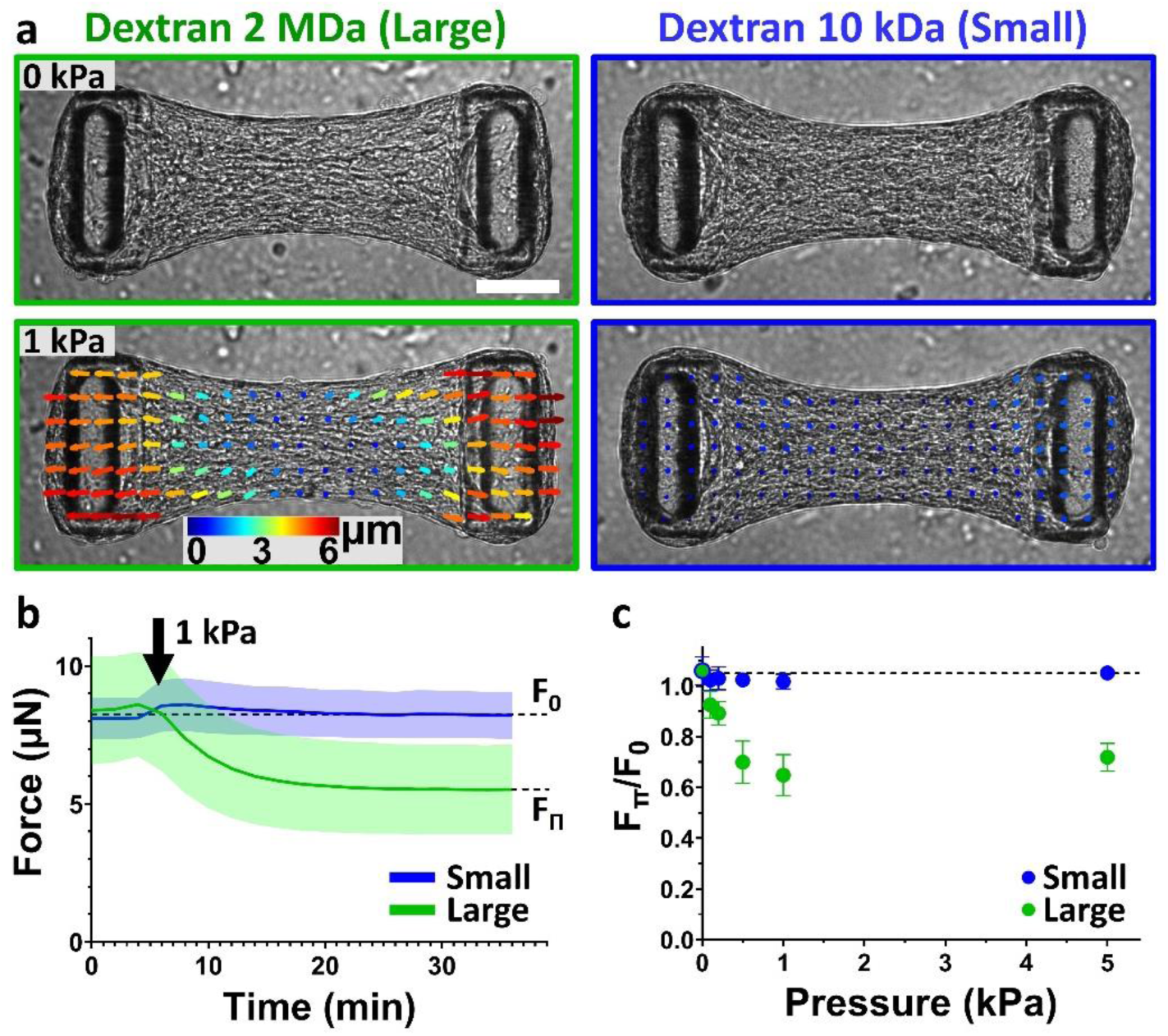
Microtissues uniaxially elongate under global but not cell compression. (a) 30 min after a 1 kPa global compression using large, 2 MDa dextran molecules (left column, outlined in green), a NIH3T3 microtissue relaxes, as shown by a representative PIV-tracking of the displacements, whereas a 1 kPa cell compression using small, 10 kDa dextran molecules has no effect (right column, outlined in blue). Scale bar is 100 µm. (b) Temporal evolution of the force generated by microtissues submitted to 1 kPa compression using large (green curve) or small (blue curve) dextran osmolytes. (c) Tissue relaxation, quantified by the force at equilibrium under pressure F_Π_ normalized by the initial force F_0_, i.e. F_Π_/F_0_, in function of the applied global (green) or cell (blue) compression. In control experiments (0 kPa), fresh culture medium without dextran was added to the microtissues. Data are the average of n > 25 microtissues over 2 independent experiments ± SD.

This relaxation could either be an active response from the cells in the form of a reduced acto-myosin contractility, or the result of a passive elongation of the extracellular matrix within the tissue by the lateral osmotic compression. Confocal images of microtissues simultaneously stained for actin, collagen and DNA showed a compacted collagen core populated with fibroblasts, surrounded by a densely cellularized peripheral shell (Supp. Movie 2). Actin stress fibers, collagen fibers and nuclei were found predominantly aligned with the tissue contraction direction (i.e. along the axis between the two cantilevers, Supp. Movie 2), consistent with fibroblasts aligning and remodeling their extracellular matrix to align with the principal maximal strains developed during tissue formation [25, 29, 39, 40].

This anisotropic architecture was previously shown to correlate with anisotropic mechanical properties, as well as anisotropic contractility [28, 33]. Based on this architecture, we propose the hypothesis that the osmotic pressure-induced elongation is not caused by passive volume-conserving effects stemming from the transient lateral tissue compression, but rather it is caused by a relaxation of the active, acto-myosin generated forces as a result of tissue compaction.

We first tested whether cells were directly sensitive to osmotic pressure by applying a pressure Π = 1 kPa selectively on the cells only using small dextran. We also observed a transient compaction of the tissue in the transverse direction (perpendicular to the long axis of the tissue), but in contrast to the response to large dextran, this transient compaction was faster and was not followed by any tissue elongation or force relaxation (F_0_ = F_Π_ = 8.1 ± 0.7 µN) (Fig. 2a-b and Supp. Movie 3). This result is consistent with the notion that cells are not directly sensitive to osmotic compression, but rather they are sensitive to the mechanical compression of the tissue, which induces active relaxation of the cells when compressed.

To test this hypothesis further, we measured the dependence of tissue relaxation in response to different pressure levels. By varying dextran concentration, we applied a compressive stress ranging from 0.1 to 5.0 kPa to microtissues while monitoring tissue contractility. This stress range corresponds to pathophysiological tissue stress induced by hydrostatic pressure [41], confined tumor growth [10, 42] or active compression by CAFs surrounding cancer cells [9]. With large dextran, force relaxation became more pronounced with increasing pressure up to 1 kPa and then plateaued (Fig. 2c). By contrast, we did not detect any significant changes in force when applying cell compression with small dextran, regardless of pressure (Fig. 2c). Together, these results point again towards an active relaxation of cell contractility in response to compression, as passive tissue elongation by the action of increasing compressive stress would be expected to monotonically increase. Also, these results confirm that force relaxation is not caused by osmotic pressure applied directly to the cells but requires the pressure to act at the tissue level.

In order to delineate the respective roles of cell and matrix within the tissue, we next investigated the impact of pressure on isolated cells and decellularized matrix.

### Without each other, neither cells nor matrix relax under pressure

To assess the contractile response of single cells to osmotic pressure, we quantified the forces generated by single fibroblasts spread on a soft PDMS substrate (Young’s modulus E = 15 kPa) grafted with 0.2 µm diameter fluorescent beads as fiducial markers and coated with fibronectin. Of note, PDMS was used instead of polyacrylamide hydrogel as the latter would be compressed by osmotic pressure, whereas PDMS is not. Cell forces and overall contractile elastic energy was calculated from substrate deformation, induced by cell-generated forces, determined from analysis of fluorescent bead images before and after cell removal [43–46]. We found that cell morphology, force localization and overall contractile energy E_C_ were unaffected by the application of a 1 kPa osmotic pressure using either small or large dextran (Fig. 3a-b), confirming that cells are not directly sensitive to osmotic pressure at that level.

**Fig. 3.**
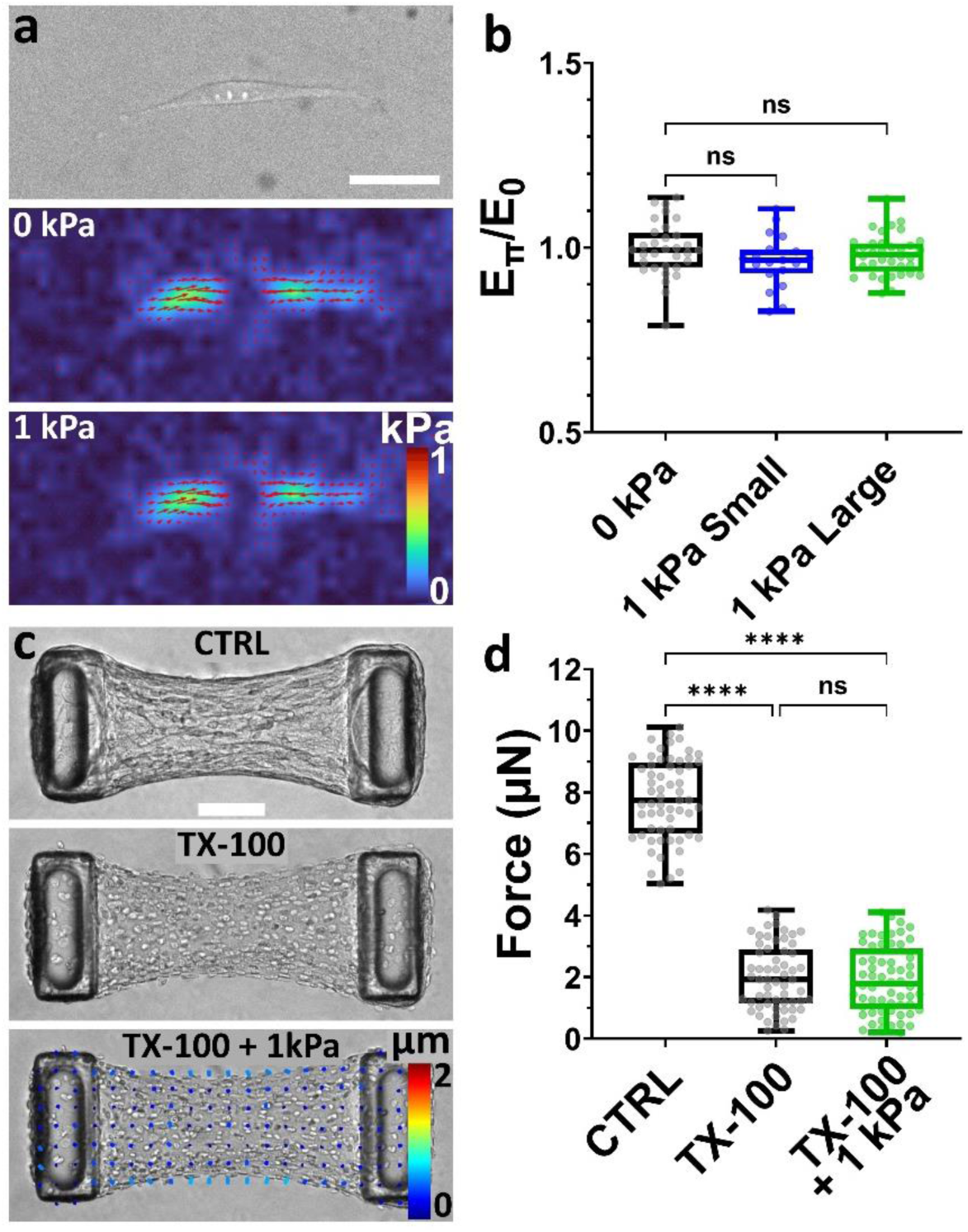
Single cells or decellularized matrix do not relax under osmotic pressure. (a) Representative brightfield image and corresponding stress maps of a NIH3T3 fibroblast spread on a soft PDMS substrate before and after application of a 1 kPa pressure using large, 2MDa dextran. Scale bar is 50 µm. (b) Contractile energy exerted by fibroblasts on the substrate under pressure (E_Π_) applied with either small (in blue) or large (in green) dextran, normalized by the initial contractile energy E_0_. In control experiments (0 kPa, in black), fresh culture medium without dextran was added to the cells. Data are presented as box plots superimposed with a dot plot of the data distribution with n > 19 cells over 2 independent experiments. (c) Representative brightfield images of a NIH3T3 microtissue (CTRL), decellularized using 0.5 % Triton X-100 (TX-100), and 30 min after the application, once decellularized, of a 1 kPa pressure using large, 2 MDa dextran (TX-100 + 1 kPa). The bottom image is superimposed with a PIV-tracking of the displacements. Scale bar is 100 µm. (d) Corresponding tissue force. Data are presented as box plots superimposed with a dot plot of the data distribution with n = 40 microtissues over 2 independent experiments. n.s. stands for non-significant (i.e. P > 0.05). ****P < 0.0001 between conditions.

To elucidate the contribution of the collagen matrix, we decellularized the microtissue using the surfactant Triton X-100. Such decellularization process was previously shown not to affect microtissue stiffness and collagen integrity [17, 47]. Contractility was drastically reduced upon decellularization, however the forces measured by pillar deflection did not fall to zero, likely because the collagen matrix was irreversibly compacted and cross-linked during tissue formation and maturation [17, 33]. We then submitted decellularized microtissues to an osmotic pressure using large, 2MDa dextran. We observed a slight compression of the tissue indicating that cell-compacted collagen laden with dead cells is not permeable to large dextran molecules, and is therefore compressed by osmotic pressure. However, we did not measure any significant displacement of the pillar tips and hence no force changes (Fig. 3.c-d and Supp. Movie 4), confirming that the elongation of intact microtissues is not driven by a passive mechanical response of the collagen under compression. This interpretation is further supported by our observation that the collagen density did not influence the relaxation of microtissues submitted to osmotic pressure (Supp. Fig. 2).

Overall, these results demonstrate that neither the cells alone nor the matrix alone relax under weak osmotic pressure. Rather, the osmotic pressure-induced elongation of microtissues is caused by the relaxation of active cell forces, in response to a mechanical compression of their surrounding tissue

### Pressure-induced relaxation depends on cell contractility

To probe the response of different tissues on pressure-induced relaxation, we applied a 1 kPa osmotic pressure on microtissues composed of different cell types known to exert different levels of contractility: murine colon carcinoma CT26 cells, murine NIH3T3 fibroblasts and human primary cancer-associated fibroblasts (CAFs) (Fig. 4.a). After 24 h of formation, CT26 microtissues reached an initial force F_0_ of 3.4 ± 2.0 µN while 3T3 microtissues generated 9.3 ± 2.4 µN (Fig. 4.b). Because primary human CAFs present a myofibroblastic phenotype [48, 49], they are larger and considerably more contractile than CT26 or 3T3 cells [50]. Consequently, we halved the cell density in CAF-based microtissues, to prevent the tissue from tearing or slipping off the cantilever. They still produced an initial force F_0_ of 25.4 ± 4.5 µN after 24 h of formation (Fig. 4.b).

**Fig. 4.**
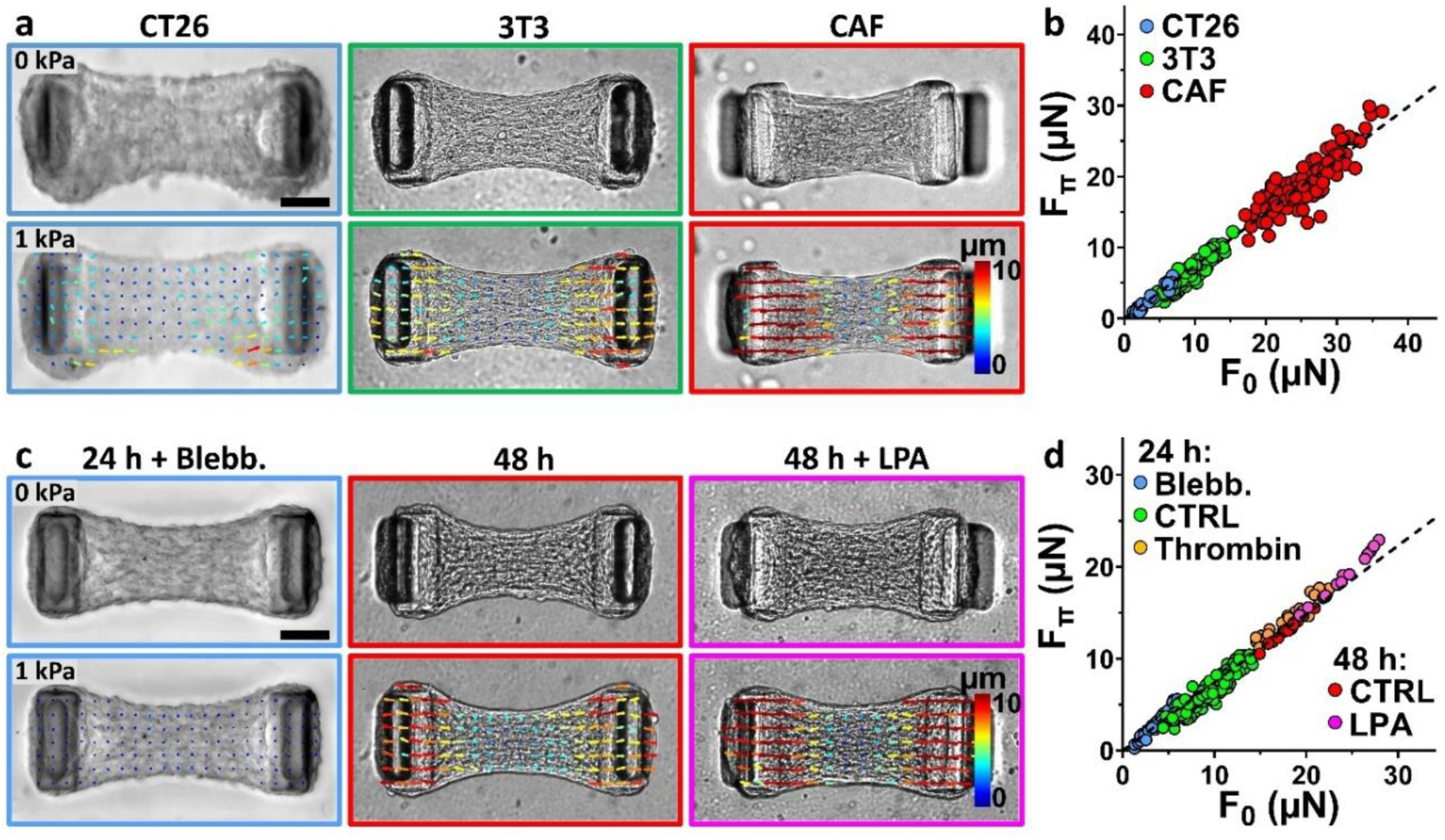
Pressure-induced relaxation correlates with the initial tissue force. (a) Representative brightfield images of microtissues composed of murine colon carcinoma CT26 cells (blue frame), murine NIH3T3 fibroblasts (green frame) or human primary cancer-associated fibroblasts (CAF, red frame) before (top row) and after (bottom row) the application of a 1 kPa pressure using large, 2 MDa dextran. The bottom image is superimposed with a PIV-tracking of the displacements. (b) Corresponding scatter plot of the pressured (F_Π_) versus the initial (F_0_) tissue force for the three cell types, superimposed with a linear regression of the data (slope = 0.74, R² = 0.97, Pearson coefficient r = 0.98). (c) Representative brightfield images of 3T3 microtissues after 24 h of formation and incubated for 1 h with 10 µM of blebbistatin (blue frame), after 48 h of formation (red frame) or after 48 h of formation and incubated for 1 h with 10 µM of lysophosphatidic acid (48 h + LPA, magenta frame) before (top row) and after (bottom row) the application of a 1 kPa pressure using large, 2 MDa dextran. The bottom image is superimposed with a PIV-tracking of the displacements. (d) Scatter plot of the pressured (F_Π_) versus the initial (F_0_) tissue force for 3T3 microtissues either after 24 h of formation and incubated for 1 h with 10 µM of blebbistatin (Blebb., blue dots), growth medium (CTRL, green dots) or 1 U/mL of thrombin (Thrombin, orange dots); or after 48 h of formation and incubated with growth medium (CTRL, red dots) or 10 µM of lysophosphatidic acid (LPA, magenta dots). The dot plot is superimposed with a linear regression of the data (slope = 0.74, R² = 0.97, Pearson coefficient r = 0.99). Scale bars are 100 µm.

Upon osmotic pressure application, all three types of microtissues relaxed, suggesting that pressure-induced relaxation is not tissue- or cell-type specific. However, the amplitude of the force relaxation linearly increased with the contractile force: weakly contractile CT26 microtissues relaxed by 1.0 ± 0.5 µN, 3T3 microtissues by 3.0 ± 0.8 µN and highly contractile CAF microtissues by 6.2 ± 2.3 µN, with a cell type-independent 26% relaxation of the initial force under pressure (Fig. 4.b). Put differently, the final force after pressure application (F_Π_) reached 74% of the initial force (F_0_), in all tissues and regardless of cell type: F_Π_ = 0.74 F_0_.

We next tested if this relationship between initial and final force also holds within a single tissue when acto-myosin-generated forces are pharmacologically altered. We cultured 3T3 microtissues for 24 h and 48 h, as maturation duration has been shown to strongly impact tissue contractility [17, 31, 33], and incubated the tissues for 1 h with either growth medium (control condition), blebbistatin (a myosin ATPase inhibitor that lowers cell contractility [31, 51, 52]), lysophosphatidic acid (LPA, a phospholipid stimulant of myosin activity via RhoA activation, which increases cell contractility [53–56]) or thrombin (an enzyme increasing RhoA-mediated myosin-generated contraction [55, 57–60]) (Fig. 4c and Supp. Fig. 3). Depending on treatment, we obtained 3T3 microtissues with initial forces F_0_ ranging from 3.5 ± 1.2 µN (24 h of maturation + 10 µM Blebbistatin) to 23.6 ± 2.8 µN (48 h of maturation + 10 µM of LPA). When subjected to an osmotic pressure of 1 kPa, all microtissues, regardless of initial force, relaxed on average by 26% (Fig. 4.c-d and Supp. Fig. 3), confirming that the pressure-induced force relaxation is regulated by myosin activity.

## Discussion

Cells constantly generate forces to migrate, to communicate mutual positions, to rearrange the surrounding ECM and neighboring cells, while also reacting to their environment, resulting in a complex feedback loop. This interplay between internal and external forces is especially crucial for tumor progression [2, 7, 9, 10]. However, the impact of cell proliferation-induced compression on cell contractility, which in turn regulates cell proliferation, remains elusive.

Here we applied osmotic compression to 3D microtissues composed of cells and collagen I while simultaneously measuring microtissue shape and contractility. In usual tensional probes, cells rapidly reinforce their cytoskeleton and increase the forces they generate in response to increased external forces [1, 5, 61, 62]. Conversely, we observed a fast and significant relaxation of microtissues subjected to a global osmotic compression and the amplitude of relaxation was correlated to the level of the imposed osmotic pressure.

We first tested whether this relaxation was an active, cell-driven mechanism, or a passive, collagen-driven one. We applied selective compression on cells only using small dextran molecules, capable of permeating between the cells and through the collagen network, but we did not measure any relaxation. These results suggest that the collagen matrix acts as a pressure sensor for cells, in agreement with our previous work showing that global compressions regulate spheroid growth and cell motility, whereas selective compression of cells only has no effect [19].

In the literature it is reported that large osmotic and mechanical pressures (of the order of hundreds of kPa) are needed to induce cell and nucleus deformation [63, 64] and interfere with cell proliferation [65–68]. The osmotic pressures we applied, ranging from 0.1 to 5 kPa, were proved to be too weak to induce measurable single cell or nucleus deformation [35]. However, cell-compacted collagen, whose elastic modulus is of the order of a few tens of kPa [17, 20, 33, 69], could be deformed by osmotic pressures in the kPa range, i.e. close to solid stress existing in tumors tissues [6, 10] .

By quantifying the forces generated by single cells spread on 2D PDMS substrates, we confirmed that cell shape and contractility were not affected by osmotic pressure in the weak compression regime we adopt. To investigate the mechanical behavior of the collagen component only, we decellularized already assembled microtissues to obtain passive collagen constructs containing dead cells. We thus observed a lack of relaxation under osmotic pressure, confirming that collagen is not directly elongating under pressure.

If cells respond to a small deformation of collagen by decreasing their contractility, we hypothesized that this response should depend on the initial contractility of the tissue, as the mechanical properties of collagen are highly dependent on its stress state [20, 33, 69, 70]. Using three different cell types as well as different drugs affecting myosin activity, we demonstrated that pressure-induced relaxation is intrinsic to cell/collagen-based constructs and is proportional to the tissue’s initial contractility, i.e. a force under a pressure of 1 kPa is always around 26 % lower than the initial force.

We speculate that collagen compression may lead to changes in ligand density, pore size or network stiffness [20, 70] that have been shown to affect cell forces [70, 71].

Overall, our results evidence a general, robust and rapid mechanosensitive response of tissues to osmotic pressure, driven by myosin activity. This mechanism could be similar to the stretch-induced fluidization of 2D cell cultures [72, 73], 3D cell/collagen-based microtissues [27] or ex vivo tissue strips [74], where external stretch leads to actin depolymerization, resulting in cell or tissue softening accompanied by a decrease in contractility [27, 72, 74]. An alternative hypothesis would be that osmotic pressure may lead to buckling of collagen or actin fibers [63, 75, 76], which was previously shown to induce drastic softening of collagen [70, 77] or severing and depolymerization of actin [75, 78], respectively.

This mechanosensitive response potentially underlies the growth inhibition previously demonstrated in tumors under compression [9, 11, 15, 16, 19], as cell contractility has been shown to regulate cell proliferation [21–24]. Thus, a thorough characterization of the mechanics of cell-compacted collagen networks under compression could be key to furthering our understanding of the relationship between compression, tissue mechanics and cell contractility. Such characterization could also put into perspective our current knowledge of volume change and stress distribution in tumor spheroids under self- or externally-generated pressure [9, 19].

In conclusion, as cell shape and contractility are key regulators of cell proliferation, epithelial-mesenchymal transition and differentiation [7, 21, 79–81], our approach combining microtissue engineering, real-time tissue force monitoring and selective osmotic pressure could pave the way to test whether the active decrease of cell contractility under pressure is a key mechanism during onco- and morphogenesis.

## Materials and Methods

### Cell culture and reagents

Mouse colon adenocarcinoma CT26 cells (ATCC CRL-2638) and NIH 3T3 fibroblasts (ATCC CRL-1658) were cultured (< 15 passages) in culture medium composed of Dulbecco’s modified Eagle’s medium (DMEM, Gibco Invitrogen) supplemented with 10% (v/v) fetal bovine serum (FBS, Gibco Invitrogen), 100 U/ml of penicillin and 100 µg/ml of streptomycin (Gibco Invitrogen). Human primary cancer-associated fibroblasts (CAF) were kindly provided by D. Matic Vignjevic (I. Curie, Paris). They were isolated from an upper rectum lieberkuhnian adenocarcinoma and immortalized as described earlier [9] at Institut Curie Hospital, Paris, with the patient’s written consent and approval of the local ethics committee. CAFs were cultured in the same culture medium supplemented with 1% (v/v) Insulin-Transferrin-Selenium (ITS, Sigma). All cell types were kept at 37°C in an atmosphere saturated in humidity and containing 5% CO2.

Triton X-100 (Sigma), Blebbistatin (Sigma), lysophosphatidic acid (Sigma) and thrombin (ThermoFisher Scientific) were introduced in the culture medium 1 h prior to the application of osmotic pressure at a 0.5% (v/v), 10 μM, 10µM and 1U/mL concentration, respectively.

### Device fabrication, calibration and microtissue engineering

The microtissues were engineered within polydimethylsiloxane (PDMS, Sylgard 184, Dow-Corning) microwells 800 µm long, 400 µm wide and 200 µm deep. Each microwell contains two T-shape cantilevers that constrain self-assembly and ensure good anchorage of the microtissue. Microwells were made by PDMS replication of SU-8-based masters microfabricated as described previously [31–33]. Briefly, successive layers of negative and positive photoresist (Microchem) were spin coated, insolated and baked to create multilayers templates. PDMS microwells were then molded from the SU-8-based masters by double replication after silanization with trichloro(1H,1H,2H,2H-perfluorooctyl)silane (Sigma) to facilitate subsequent release of PDMS from the template. PDMS stiffness was assessed through uniaxial tensile tests with an Instron 5848 Microtester (Instron). Cantilever spring constant *k* was calibrated with a capacitive MEMS force sensor mounted on a micromanipulator as described previously [29, 31, 33, 82] and found to be *k* = 0.45 ± 0.10 N/m.

PDMS microwells were sterilized in 70 % ethanol and treated with 0.2 % (m/v) Pluronic F127 (Sigma) for 2 min to reduce cell adhesion. A cooled suspension of 300,000 cells (CAF) or 600,000 cells (CT26 and 3T3) per mL of liquid neutralized collagen I from rat tail (Advanced Biomatrix) was then added to the microwells on ice and centrifuged to drive cells into the recessed wells. The collagen concentration was set at 2.0 mg/mL for all experiments except for Supp. Fig. 2 where it was set at 1.5 or 2.5 mg/mL. Excess cell/collagen mixture was removed and the remaining constructs were polymerized at 37°C for 9 min before the addition of culture medium. Microtissues were kept in the incubator for 24 h or 48 h prior to experiments. Over time, cells spread and compact the collagen matrix to form a microtissue suspended between the top of the pair of cantilevers.

### Osmotic pressure

Osmotic pressure is exerted by adding to the culture medium a well-defined amount of dextran (Sigma), a biologically inert polysaccharide. As such, its concentration gradient cannot be balanced by the activity of ion pumps, channels or endocytosis and it therefore exerts a persistent osmotic stress [13, 19, 35, 36]. We used large dextran (MW = 2 MDa, hydrodynamic radius > 25 nm [38]) which cannot penetrate the cell/collagen network to apply osmotic stress to the entire microtissue (Fig. 1.b top schematic). We used small dextran (MW = 10 kDa, hydrodynamic radius < 2nm [38]) that can diffuse between cells and through collagen to apply pressure only to cells (Fig. 1.b bottom schematic). The relationships between dextran size, concentration and resulting osmotic pressure are given in Table 1.

**Table 1.** Range of osmotic pressure used and corresponding concentrations of large, 2 MDa dextran and small, 10 kDa dextran. For small, 10 kDa dextran, the osmotic pressure is directly derived from the van ’t Hoff formula Π = *c*.*R*.*T*, with *c* the molar concentration of dextran, *R* the ideal gas constant, and *T* the absolute temperature. Because of their size, 2 MDa dextran molecules are self-interacting and do not obey the van ’t Hoff equation. The relationship between concentration and pressure has been previously calibrated [35, 83–85]

| Osmotic pressure (kPa) | [2 Mda Dextran] (g/L) | [10 kda Dextran] (g/L) |
| --- | --- | --- |
| 0.1 | 3.2 | 0.4 |
| 0.2 | 5.9 | 0.8 |
| 0.5 | 12.4 | 2.0 |
| 1.0 | 20.6 | 4.0 |
| 5.0 | 54.8 | 20.0 |

### Microscopy and image analysis

Brightfield imaging was performed using either a Nikon Eclipse TI-2 inverted microscope equipped with an Orca flash 4.0 LT CMOS digital camera (Hamamatsu), a CFI S Plan Fluor ELWD 20x/0.45 objective (Nikon) and controlled with NIS Elements software (Nikon), or a Nikon Ti-E inverted microscope equipped with a Zyla sCMOS camera (Andor), a Plan Fluor10x/0.30 objective (Nikon) and controlled with IQ3.1 software (Andor). Both microscopes are equipped with an incubator that maintains an atmosphere of 37°C, saturated with humidity and containing 5% CO_2_.

The force generated by individual microtissues before, during and after osmotic pressure was assessed from the deflection of the cantilevers, as previously described [31–33]. Briefly, the deflection *d* was determined by comparing the position of the cantilevers’ T-shaped caps with their initial position (i.e. before tissue formation). Brightfield images were taken every 2 min and the position of the top of the cantilevers was tracked using a previously developed MATLAB script [33]. Tracking results were checked visually and faulty tracking was either redone by hand or discarded. The force *F* generated by the microtissue was deduced from the average deflection *d_AVG_* of the two cantilevers and spring constant *k* as follow: *F* = *k*.*d_AVG_*. Only tissues that were uniformly anchored to the tips of the two cantilevers throughout the duration of the experiments were included in the analysis.

Displacement fields were calculated from the brightfield images using a Matlab particle image velocimetry (PIV) toolbox (https://pivlab.blogspot.com/) [86], as previously described [33]. Briefly, the algorithm cross-correlates small interrogation areas of a pair of images (reference image at t = 0 and image of interest) in the frequency domain using FFT to determine the most likely displacement vector (u,v) of the particle at position (x,y) in the interrogation area. The 1500x450 pixels (660x330 µm) images were analyzed in four successive passes with decreasing interrogation areas, from 128×128 to 32×32 pixels leading to a final resolution of 14 µm. Missing vectors were replaced by interpolated data [87], outliers were filtered using a local normalized median filter [88], and the noise was reduced using a penalized least squares method [89]. It should be noted that cell migration during the duration of a pressure-induced relaxation was considered negligible compared to cell deformation. For readability reasons, only half of the vectors are represented in the displacement-superimposed images.

### Traction force microscopy (TFM) of single cells

Force measurements were performed using a method described previously [90]. In short, fluorescent beads were grafted to the surface of embedded in a soft PDMS substrate with 15 kPa rigidity and images of those beads were taken before and during the application of osmotic pressure. Soft PDMS (Dowsyl CY 52-276) substrates were prepared by mixing CyA and CyB components at 1:1 ratio before spin-coating 0.1 g for 30s at 500 rpm on a 35 mm glass bottom Fluorodish (World Precision Instruments) to achieve a flat 60 - 100 μm flat layer. After curing at 80°C for 2 hours, the substrate was silanized with 10% (3-Aminopropyl) trimethoxysilane (APTES, Sigma) in ethanol for 15 min before grafting carboxylated 200 nm fluorescent beads (647 nm, Invitrogen) at a 4:500 dilution in deionized water. Bovine fibronectin (50 μg/ml, Sigma) was incubated on the substrate for 1 hour prior to cell seeding. Between each step, the samples were rinsed 3 times with PBS. Approximately 50,000 cells were seeded and let to adhere overnight. At the end of the experiment, cells were removed using a deionized water-based osmotic shock and an unstressed reference image of the beads was taken.

The displacement field analysis was obtained using a homemade algorithm based on optical flow [91, 92]. Cellular traction forces were calculated using Fourier transform traction cytometry with zero-order regularization [91, 93, 94] under the assumption that the substrate is a linear elastic half-space and considering only displacement and stress tangential to the substrate. To calculate the strain energy stored in the substrate, the scalar product of the stress and displacement vector fields was integrated over the surface of the whole cell. The algorithm was implemented in Python and is available in [91, 92].

TFM movies were recorded on a Nikon Ti2 eclipse microscope equipped with a Yokogawa CSU-W1 spinning disk confocal scanner and a Teledyne Photometrics Prime BSI camera at 20x magnification (NA = 0.75).

### Statistics

For each box plots, the box extends from the 25th to 75th percentiles, the median is plotted as a line inside the box, the whiskers extend to the most extreme data point and the data distribution is superimposed as a dot plot. Statistical significances were determined by one-way analysis of variance (ANOVA) corrected for multiple comparisons using Tukey test with Prism (GraphPad).

## Acknowledgments

F.W. and S.d.B. acknowledge financial support from the LabEx Who Am I?, grant ANR-11-LABX-0071, and the Initiatives d’Excellence 572, grant ANR-11-IDEX-0005-02 TP5. P.M. acknowledges financial support from the Marie Skłodowska-Curie Action Individual Fellowship for the Project CHECKMATE (grant 893981). J.F. acknowledges financial support from ITMO Cancer of Aviesan within the framework of the 2021-2030 Cancer Control Strategy, on funds administered by Inserm. M.B. acknowledges financial support from the ANR PlatforMech project, grant ANR-18-CE14–0037-02 and the ANR Inter-s-cal project, grant ANR-21-CE13–0042-02 of the French Agence Nationale de la Recherche (ANR). G.C. acknowledges financial support from the ANR SupraWaves project, grant ANR-19-CE13-0028 of the French ANR. T.B. and S.d.B. acknowledge funding from CNRS grants (PEPS CNRS-INSIS 2021, Lumière Visible et Vie 2022). T.B. and B.F. acknowledge funding from the ANR CONTRACTILE project, grant ANR-23-CE13-0037 of the French ANR. The authors thank O. Zajac and D. Matic Vignjevic (I. Curie, Paris) for kindly providing the human primary cancer-associated fibroblasts, as well as D. Debarre (LIPhy, Grenoble) for help with image analysis.

## Author Contributions Statement

M.B., G.C. and T.B. conceived the study and designed the experiments. F.W., G.M-V, A.E.R., P.A., J.F., G.C. and T.B. performed experiments. G.M-V, A.E.R., P.A., P.M., B.F., D.A., P.R., G.C. and T.B. analyzed the data. G.C. and T.B. wrote the manuscript with feedback from all authors. M.B., G.C. and T.B. supervised the project.

## Data availability

Because of the large file size, the datasets generated and/or analyzed during the current study are available from the corresponding author on request. A response will be provided in less than 2 weeks.

## Supplementary Information

**Supp. Movie 1. 3T3 microtissue response to tissue compression.** Time lapse of a representative 3T3 microtissue submitted to a 1 kPa osmotic pressure using large, 2MDa dextran molecules at t = 12 min. Upon osmotic pressure, the PIV-tracking of the displacements highlights the transient, transverse compression of the tissue, followed by its elongation.

**Supp. Movie 2. Confocal reconstruction of an 3T3 microtissue.** Merged Z-stack (top) of a representative 3T3 microtissue stained for actin (in green), collagen (in magenta) and nuclei (in blue), highlighting the anisotropic orientation of actin and collagen fibers along the x-axis. The right column shows magnifications of the central region for each staining. Scale bars are 50 µm.

**Supp. Movie 3. 3T3 microtissue response to cell-only compression.** Time lapse of a representative 3T3 microtissue submitted to a 1 kPa osmotic pressure using small, 10kDa dextran at t = 12 min. Upon osmotic pressure, the PIV-tracking of the displacements highlights the transient compaction of the tissue in the transverse direction, not followed by any tissue elongation.

**Supp. Movie 4. Decellularized 3T3 microtissue response to tissue compression.** Time lapse of a representative 3T3 microtissue decellularized with 0.5 % Triton X-100 and submitted to a 1 kPa osmotic pressure using large, 2MDa dextran molecules at t = 12 min. Upon osmotic pressure, the PIV-tracking of the displacements highlights the slight compression of the tissue, not followed by any tissue elongation.

**Supp. Fig. 1.**
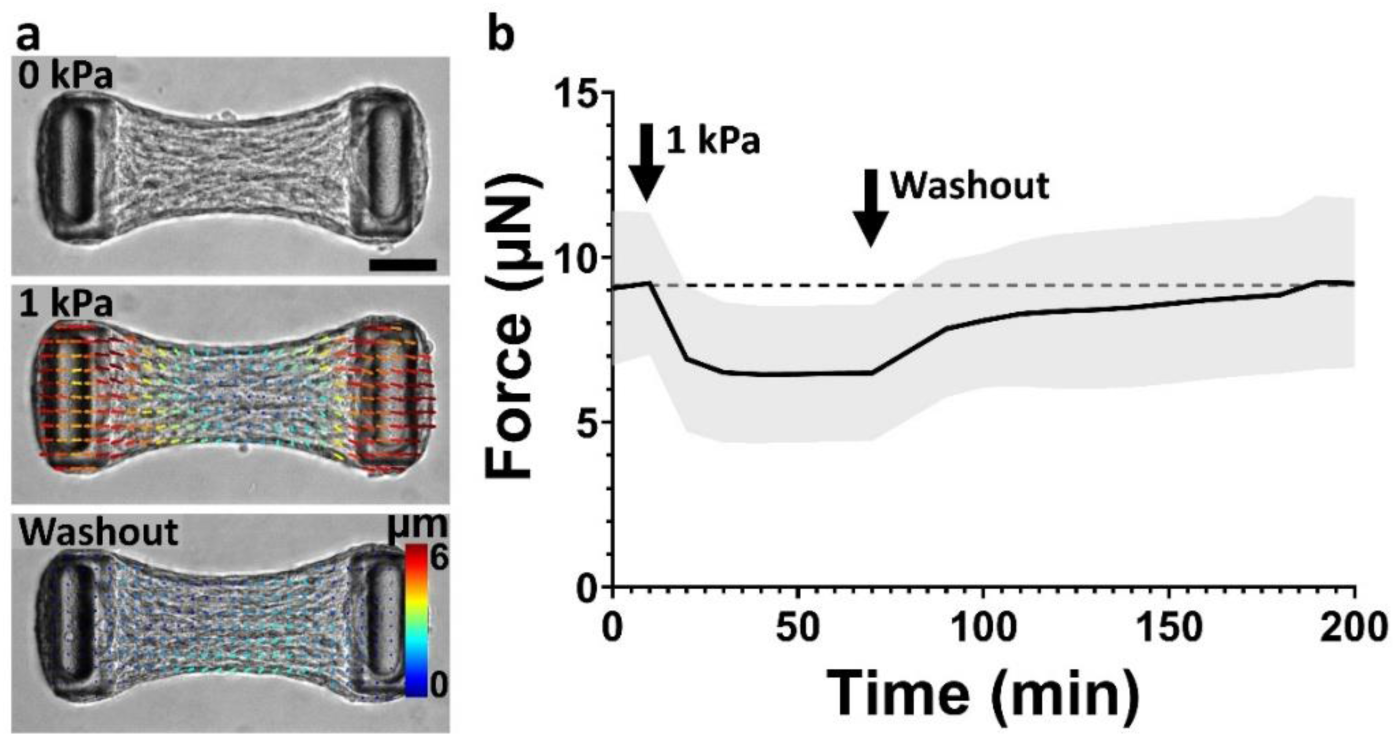
Pressure-induced relaxation is reversible. (a) PIV-tracking of the relaxation and recovery of a representative NIH3T3 microtissue upon the application and removal of a 1 kPa global compression using large, 2 MDa dextran molecules, respectively. (b) Temporal evolution of the force generated by microtissues submitted to the application and washout of a 1 kPa compression using large dextran osmolytes. Data are the average of n > 50 microtissues over 3 independent experiments ± SD. Scale bar is 100 µm.

**Supp. Fig. 2.**
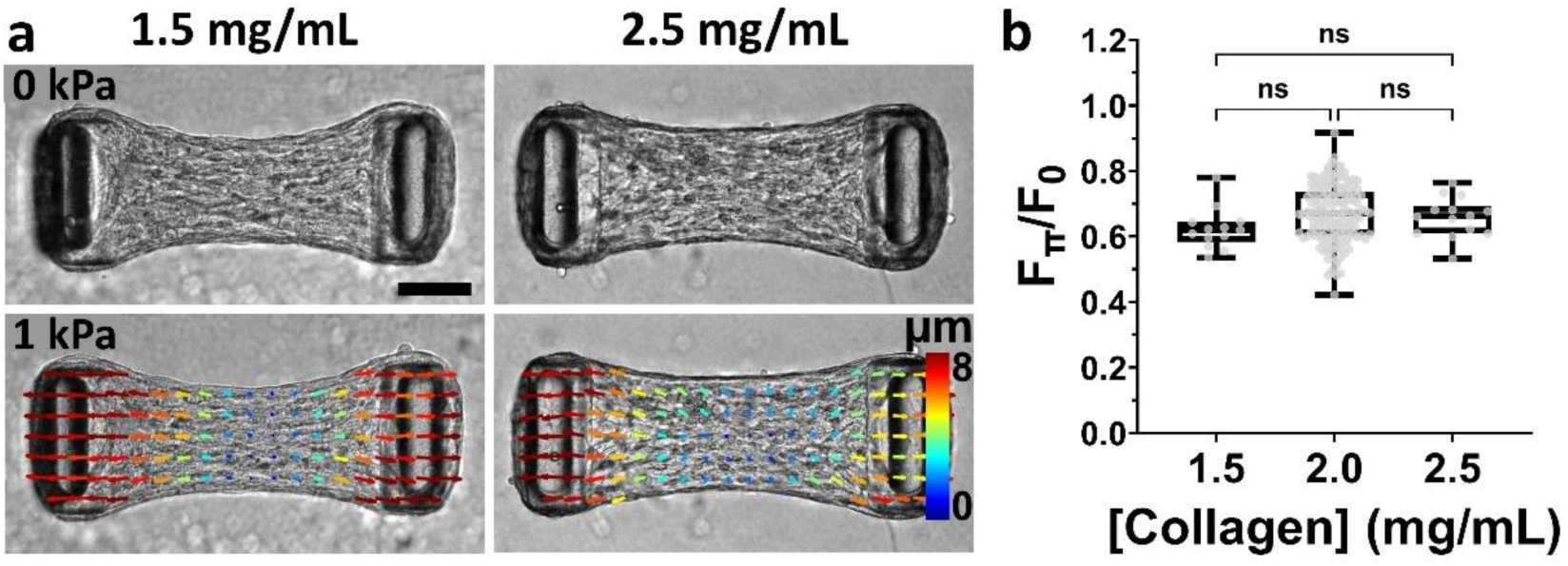
Pressure-induced relaxation is independent of initial collagen density. (a) PIV-tracking of the relaxation of representative NIH3T3 microtissues composed of 1.5 mg/mL (left column) or 2.5 mg/mL (right column) collagen upon the application of a 1 kPa global compression using large, 2 MDa dextran. (b) Tissue relaxation, quantified by the force under pressure F_Π_ normalized by the initial force F_0_, i.e. F/F_0_, as a function of the collagen concentration. Data are presented as box plots superimposed with a dot plot of the data distribution with n > 13 microtissues over 2 independent experiments. n.s. stands for non-significant (i.e. P > 0.05). Scale bar is 100 µm.

**Supp. Fig. 3.**
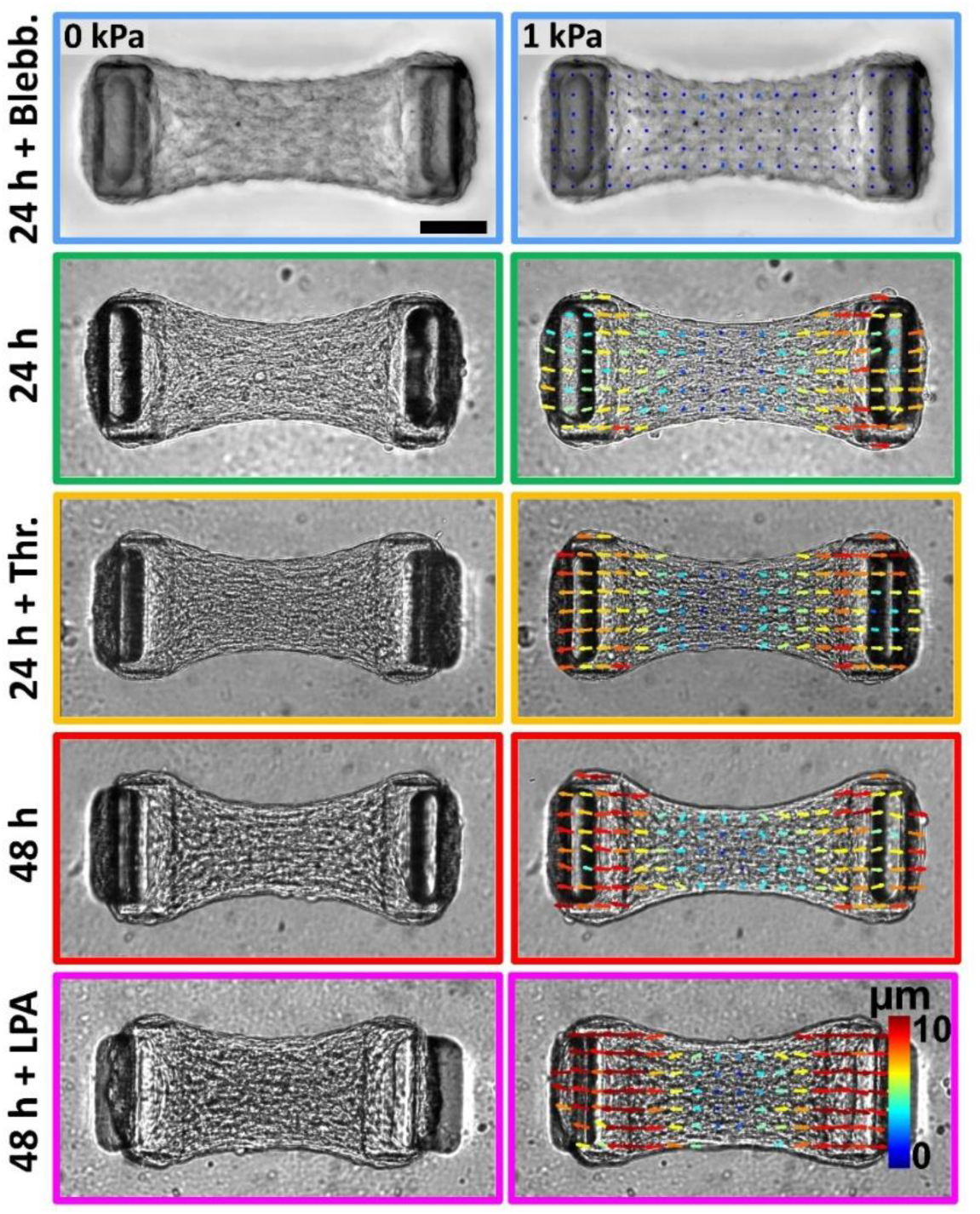
Pressure-induced relaxation correlates with the initial tissue force. (a) Representative brightfield images of 3T3 microtissues after 24 h of formation and incubated for 1 h with either 10 µM of blebbistatin (24h + Blebb., blue frame), growth medium (24 h, green frame) or 1 U/mL of thrombin (24 h + Thr.), after 48 h of formation (48 h, red frame) or after 48 h of formation and incubated for 1 h with 10 µM of lysophosphatidic acid (48 h + LPA, magenta frame) before (left column) and after (right column) the application of a 1 kPa pressure using large, 2 MDa dextran. The image in the right-hand column is superimposed with a PIV-tracking of the displacements. Relaxation quantification is shown in Fig. 4. Scale bar is 100 µm.

## References

1. Eyckmans J, Boudou T, Yu X, Chen CS (2011) A Hitchhiker’s Guide to Mechanobiology. Dev Cell 21:35–47. 10.1016/j.devcel.2011.06.015

2. Nia HT, Munn LL, Jain RK (2020) Physical traits of cancer. Science (80- ) 370:. 10.1126/SCIENCE.AAZ0868

3. Humphrey JD, Dufresne ER, Schwartz MA (2014) Mechanotransduction and extracellular matrix homeostasis. Nat Rev Mol Cell Biol 15:802–812. 10.1038/nrm3896

4. Saraswathibhatla A, Indana D, Chaudhuri O (2023) Cell–extracellular matrix mechanotransduction in 3D. Nat Rev Mol Cell Biol 24:495–516. 10.1038/s41580-023-00583-1

5. Romani P, Valcarcel-Jimenez L, Frezza C, Dupont S (2021) Crosstalk between mechanotransduction and metabolism. Nat Rev Mol Cell Biol 22:22–38. 10.1038/s41580-020-00306-w

6. Nia HT, Liu H, Seano G, Datta M, Jones D, Rahbari N, Incio J, Chauhan VP, Jung K, Martin JD, Askoxylakis V, Padera TP, Fukumura D, Boucher Y, Hornicek FJ, Grodzinsky AJ, Baish JW, Munn LL, Jain RK (2016) Solid stress and elastic energy as measures of tumour mechanopathology. Nat Biomed Eng 1:0004. 10.1038/s41551-016-0004

7. Broders-Bondon F, Ho-Bouldoires THN, Fernandez-Sanchez ME, Farge E (2018) Mechanotransduction in tumor progression: The dark side of the force. J Cell Biol 217:1571– 1587. 10.1083/jcb.201701039

8. Levental KR, Yu H, Kass L, Lakins JN, Egeblad M, Erler JT, Fong SFT, Csiszar K, Giaccia A, Weninger W, Yamauchi M, Gasser DL, Weaver VM (2009) Matrix Crosslinking Forces Tumor Progression by Enhancing Integrin Signaling. Cell 139:891–906. 10.1016/j.cell.2009.10.027

9. Barbazan J, Pérez-González C, Gómez-González M, Dedenon M, Richon S, Latorre E, Serra M, Mariani P, Descroix S, Sens P, Trepat X, Vignjevic DM (2023) Cancer-associated fibroblasts actively compress cancer cells and modulate mechanotransduction. Nat Commun 14:. 10.1038/s41467-023-42382-4

10. Stylianopoulos T, Martin JD, Chauhan VP, Jain SR, Diop-Frimpong B, Bardeesy N, Smith BL, Ferrone CR, Hornicek FJ, Boucher Y, Munn LL, Jain RK (2012) Causes, consequences, and remedies for growth-induced solid stress in murine and human tumors. Proc Natl Acad Sci U S A 109:15101–15108. 10.1073/pnas.1213353109

11. Helmlinger G, Netti PA, Lichtenbeld HC, Melder RJ, Jain RK (1997) Solid stress inhibits the growth of multicellular tumor spheroids. Nat Biotechnol 15:778–783. 10.1038/nbt0897-778

12. Alessandri K, Sarangi BR, Gurchenkov VV, Sinha B, Kießling TR, Fetler L, Rico F, Scheuring S, Lamaze C, Simon A, Geraldo S, Vignjević D, Doméjean H, Rolland L, Funfak A, Bibette J, Bremond N, Nassoy P (2013) Cellular capsules as a tool for multicellular spheroid production and for investigating the mechanics of tumor progression in vitro. Proc Natl Acad Sci U S A 110:14843–14848. 10.1073/pnas.1309482110

13. Montel F, Delarue M, Elgeti J, Malaquin L, Basan M, Risler T, Cabane B, Vignjevic D, Prost J, Cappello G, Joanny J-F (2011) Stress Clamp Experiments on Multicellular Tumor Spheroids. Phys Rev Lett 107:188102. 10.1103/PhysRevLett.107.188102

14. Delarue M, Montel F, Vignjevic D, Prost J, Joanny JF, Cappello G (2014) Compressive stress inhibits proliferation in tumor spheroids through a volume limitation. Biophys J 107:1821– 1828. 10.1016/j.bpj.2014.08.031

15. Taubenberger A V., Girardo S, Träber N, Fischer-Friedrich E, Kräter M, Wagner K, Kurth T, Richter I, Haller B, Binner M, Hahn D, Freudenberg U, Werner C, Guck J (2019) 3D Microenvironment Stiffness Regulates Tumor Spheroid Growth and Mechanics via p21 and ROCK. Adv Biosyst 3:1–46. 10.1002/adbi.201900128

16. Garau Paganella L, Badolato A, Labouesse C, Fischer G, Sänger CS, Kourouklis A, Giampietro C, Werner S, Mazza E, Tibbitt MW (2024) Variations in fluid chemical potential induce fibroblast mechano-response in 3D hydrogels. Biomater Adv 163:213933. 10.1016/j.bioadv.2024.213933

17. Zhao R, Boudou T, Wang WG, Chen CS, Reich DH (2013) Decoupling cell and matrix mechanics in engineered microtissues using magnetically actuated microcantilevers. Adv Mater 25:1699– 1705. 10.1002/adma.201203585

18. Smith ML, Gourdon D, Little WC, Kubow KE, Eguiluz RA, Luna-Morris S, Vogel V (2007) Force-induced unfolding of fibronectin in the extracellular matrix of living cells. PLoS Biol 5:2243– 2254. 10.1371/journal.pbio.0050268

19. Dolega ME, Monnier S, Brunel B, Joanny JF, Recho P, Cappello G (2021) Extra-cellular matrix in multicellular aggregates acts as a pressure sensor controlling cell proliferation and motility. Elife 10:1–33. 10.7554/eLife.63258

20. Licup AJ, Münster S, Sharma A, Sheinman M, Jawerth LM, Fabry B, Weitz DA, MacKintosh FC (2015) Stress controls the mechanics of collagen networks. Proc Natl Acad Sci U S A 112:9573– 9578. 10.1073/pnas.1504258112

21. Nelson CM, Jean RP, Tan JL, Liu WF, Sniadecki NJ, Spector AA, Chen CS (2005) Emergent patterns of growth controlled by multicellular form and mechanics. Proc Natl Acad Sci U S A 102:11594–11599. 10.1073/pnas.0502575102

22. Civelekoglu-Scholey G, Scholey JM (2010) Mitotic force generators and chromosome segregation. Cell Mol Life Sci 67:2231–2250. 10.1007/s00018-010-0326-6

23. 23. Dupont S, Morsut L, Aragona M, Enzo E, Giulitti S, Cordenonsi M, Zanconato F, Le Digabel J, Forcato M, Bicciato S, Elvassore N, Piccolo S (2011) Role of YAP/TAZ in mechanotransduction. Nature 474:179–184. 10.1038/nature10137

24. Lafaurie-Janvore J, Maiuri P, Wang I, Pinot M, Manneville J-B, Betz T, Balland M, Piel M (2013) ESCRT-III Assembly and Cytokinetic Abscission Are Induced by Tension Release in the Intercellular Bridge. Science (80- ) 339:1625–1629. 10.1126/science.1233866

25. Zhao R, Chen CS, Reich DH (2014) Force-driven evolution of mesoscale structure in engineered 3D microtissues and the modulation of tissue stiffening. Biomaterials 35:5056–5064. 10.1016/j.biomaterials.2014.02.020

26. Walker M, Godin M, Pelling AE (2018) A vacuum-actuated microtissue stretcher for long-term exposure to oscillatory strain within a 3D matrix. Biomed Microdevices 20:. 10.1007/s10544-018-0286-4

27. Walker M, Rizzuto P, Godin M, Pelling AE (2020) Structural and mechanical remodeling of the cytoskeleton maintains tensional homeostasis in 3D microtissues under acute dynamic stretch. Sci Rep 10:1–16. 10.1038/s41598-020-64725-7

28. Bose P, Eyckmans J, Nguyen TD, Chen CS, Reich DH (2019) Effects of Geometry on the Mechanics and Alignment of Three-Dimensional Engineered Microtissues. ACS Biomater Sci Eng 5:3843–3855. 10.1021/acsbiomaterials.8b01183

29. Legant WR, Chen CS, Vogel V (2012) Force-induced fibronectin assembly and matrix remodeling in a 3D microtissue model of tissue morphogenesis. Integr Biol 4:1164. 10.1039/c2ib20059g

30. Sakar MS, Eyckmans J, Pieters R, Eberli D, Nelson BJ, Chen CS (2016) Cellular forces and matrix assembly coordinate fibrous tissue repair. Nat Commun 7:1–8. 10.1038/ncomms11036

31. Legant WR, Pathak A, Yang MT, Deshpande VS, McMeeking RM, Chen CS (2009) Microfabricated tissue gauges to measure and manipulate forces from 3D microtissues. Proc Natl Acad Sci U S A 106:10097–10102. 10.1073/pnas.0900174106

32. Ramade A, Legant WR, Picart C, Chen CS, Boudou T (2014) Microfabrication of a platform to measure and manipulate the mechanics of engineered microtissues. Methods Cell Biol 121:191–211. 10.1016/B978-0-12-800281-0.00013-0

33. Méry A, Ruppel A, Revilloud J, Balland M, Cappello G, Boudou T (2023) Light-driven biological actuators to probe the rheology of 3D microtissues. Nat Commun 14:1–12. 10.1038/s41467-023-36371-w

34. Dolega M, Zurlo G, Goff M Le, Greda M, Verdier C, Joanny JF, Cappello G, Recho P (2021) Mechanical behavior of multi-cellular spheroids under osmotic compression. J Mech Phys Solids 147:104205. 10.1016/j.jmps.2020.104205

35. Monnier S, Delarue M, Brunel B, Dolega ME, Delon A, Cappello G (2016) Effect of an osmotic stress on multicellular aggregates. Methods 94:114–119. 10.1016/j.ymeth.2015.07.009

36. Joyce K, Fabra GT, Bozkurt Y, Pandit A (2021) Bioactive potential of natural biomaterials: identification, retention and assessment of biological properties. Signal Transduct Target Ther 6:1–28. 10.1038/s41392-021-00512-8

37. Dolega ME, Delarue M, Ingremeau F, Prost J, Delon A, Cappello G (2017) Cell-like pressure sensors reveal increase of mechanical stress towards the core of multicellular spheroids under compression. Nat Commun 8:1–9. 10.1038/ncomms14056

38. Armstrong JK, Wenby RB, Meiselman HJ, Fisher TC (2004) The hydrodynamic radii of macromolecules and their effect on red blood cell aggregation. Biophys J 87:4259–4270. 10.1529/biophysj.104.047746

39. Eastwood M, Mudera VC, McGrouther DA, Brown RA (1998) Effect of precise mechanical loading on fibroblast populated collagen lattices: Morphological changes. Cell Motil Cytoskeleton 40:13–21. 10.1002/(SICI)1097-0169(1998)40:1<13::AID-CM2>3.0.CO;2-G

40. Sander EA, Barocas VH, Tranquillo RT (2011) Initial fiber alignment pattern alters extracellular matrix synthesis in fibroblast-populated fibrin gel cruciforms and correlates with predicted tension. Ann Biomed Eng 39:714–729. 10.1007/s10439-010-0192-2

41. Chugh M, Munjal A, Megason SG (2022) Hydrostatic pressure as a driver of cell and tissue morphogenesis. Semin Cell Dev Biol 131:134–145. 10.1016/j.semcdb.2022.04.021

42. Fernandez-Sanchez ME, Barbier S, Whitehead J, Bealle G, Michel A, Latorre-Ossa H, Rey C, Fouassier L, Claperon A, Brulle L, Girard E, Servant N, Rio-Frio T, Marie HLN, Lesieur S, Housset C, Gennisson JL, Tanter ML, Menager C, Fre S, Robine S, Farge E (2015) Mechanical induction of the tumorigenic b-catenin pathway by tumour growth pressure. Nature 523:92–95. 10.1038/NATURE14329

43. Dembo M, Wang Y-L (1999) Stresses at the Cell-to-Substrate Interface during Locomotion of Fibroblasts. Biophys J 76:2307–2316. 10.1016/S0006-3495(99)77386-8

44. Balaban NQ, Schwarz US, Riveline D, Goichberg P, Tzur G, Sabanay I, Mahalu D, Safran S, Bershadsky A, Addadi L, Geiger B (2001) Force and focal adhesion assembly: A close relationship studied using elastic micropatterned substrates. Nat Cell Biol 3:466–472. 10.1038/35074532

45. Butler JP, Toli-Nørrelykke IM, Fabry B, Fredberg JJ (2002) Traction fields, moments, and strain energy that cells exert on their surroundings. Am J Physiol - Cell Physiol 282:595–605. 10.1152/ajpcell.00270.2001

46. Andersen T, Wörthmüller D, Probst D, Wang I, Moreau P, Fitzpatrick V, Boudou T, Schwarz US, Balland M (2023) Cell size and actin architecture determine force generation in optogenetically activated cells. Biophys J 122:684–696. 10.1016/j.bpj.2023.01.011

47. Crapo PM, Gilbert TW, Badylak SF (2011) An overview of tissue and whole organ decellularization processes. Biomaterials 32:3233–3243. 10.1016/j.biomaterials.2011.01.057

48. Santi A, Kugeratski FG, Zanivan S (2018) Cancer Associated Fibroblasts: The Architects of Stroma Remodeling. Proteomics 18:1–15. 10.1002/pmic.201700167

49. Barbazán J, Matic Vignjevic D (2019) Cancer associated fibroblasts: is the force the path to the dark side? Curr Opin Cell Biol 56:71–79. 10.1016/j.ceb.2018.09.002

50. Erdogan B, Ao M, White LM, Means AL, Brewer BM, Yang L, Washington MK, Shi C, Franco OE, Weaver AM, Hayward SW, Li D, Webb DJ (2017) Cancer-associated fibroblasts promote directional cancer cell migration by aligning fibronectin. J Cell Biol 216:3799–3816. 10.1083/jcb.201704053

51. Straight AF, Cheung A, Limouze J, Chen I, Westwood NJ, Sellers JR, Mitchison TJ (2003) Dissecting temporal and spatial control of cytokinesis with a myosin II inhibitor. Science (80- ) 299:1743–1747. 10.1126/science.1081412

52. Paszek MJ, Zahir N, Johnson KR, Lakins JN, Rozenberg GI, Gefen A, Reinhart-King CA, Margulies SS, Dembo M, Boettiger D, Hammer DA, Weaver VM (2005) Tensional homeostasis and the malignant phenotype. Cancer Cell 8:241–254. 10.1016/j.ccr.2005.08.010

53. Moolenaar WH (1995) Lysophosphatidic acid, a multifunctional phospholipid messenger. J Biol Chem 270:12949–12952. 10.1074/jbc.270.22.12949

54. Kranenburg O, Poland M, Gebbink M, Oomen L, Moolenaar WH (1997) Dissociation of LPA-induced cytoskeletal contraction from stress fiber formation by differential localization of RhoA. J Cell Sci 110:2417–2427. 10.1242/jcs.110.19.2417

55. Yang MT, Reich DH, Chen CS (2011) Measurement and analysis of traction force dynamics in response to vasoactive agonists. Integr Biol 3:663–674. 10.1039/c0ib00156b

56. 56. Geraldo LHM, Spohr TCL de S, Amaral RF do, Fonseca ACC da, Garcia C, Mendes F de A, Freitas C, dosSantos MF, Lima FRS (2021) Role of lysophosphatidic acid and its receptors in health and disease: novel therapeutic strategies. Signal Transduct Target Ther 6:. 10.1038/s41392-020-00367-5

57. Goeckeler ZM, Wysolmerski RB (1995) Myosin light chain kinase-regulated endothelial cell contraction: The relationship between isometric tension, actin polymerization, and myosin phosphorylation. J Cell Biol 130:613–627. 10.1083/jcb.130.3.613

58. Dudek SM, Garcia JGN (2001) Cytoskeletal regulation of pulmonary vascular permeability. J Appl Physiol 91:1487–1500. 10.1152/jappl.2001.91.4.1487

59. Liu Z, Tan JL, Cohen DM, Yang MT, Sniadecki NJ, Ruiz SA, Nelson CM, Chen CS (2010) Mechanical tugging force regulates the size of cell-cell junctions. Proc Natl Acad Sci U S A 107:9944–9949. 10.1073/pnas.0914547107

60. 60. Reinhard NR, Van Helden SF, Anthony EC, Yin T, Wu YI, Goedhart J, Gadella TWJ, Hordijk PL (2016) Spatiotemporal analysis of RhoA/B/C activation in primary human endothelial cells. Sci Rep 6:1–16. 10.1038/srep25502

61. Iskratsch T, Wolfenson H, Sheetz MP (2014) Appreciating force and shape-the rise of mechanotransduction in cell biology. Nat Rev Mol Cell Biol 15:825–833. 10.1038/nrm3903

62. Schwartz MA (2010) Integrins and Extracellular Matrix in Mechanotransduction. Cold Spring Harb Perspect Biol 2:a005066–a005066. 10.1101/cshperspect.a005066

63. Zhou EH, Trepat X, Park CY, Lenormand G, Oliver MN, Mijailovich SM, Hardin C, Weitz DA, Butler JP, Fredberg JJ (2009) Universal behavior of the osmotically compressed cell and its analogy to the colloidal glass transition. Proc Natl Acad Sci U S A 106:10632–10637. 10.1073/pnas.0901462106

64. Kim D-H, Li B, Si F, Philips J, Wirtz D, Sun SX (2015) Volume regulation and shape bifurcation in the cell nucleus. J Cell Sci 129:457. 10.1242/jcs.166330

65. Guo M, Pegoraro AF, Mao A, Zhou EH, Arany PR, Han Y, Burnette DT, Jensen MH, Kasza KE, Moore JR, Mackintosh FC, Fredberg JJ, Mooney DJ, Lippincott-Schwartz J, Weitz DA (2017) Cell volume change through water efflux impacts cell stiffness and stem cell fate. Proc Natl Acad Sci U S A 114:E8618–E8627. 10.1073/pnas.1705179114

66. Han YL, Pegoraro AF, Li H, Li K, Yuan Y, Xu G, Gu Z, Sun J, Hao Y, Gupta SK, Li Y, Tang W, Kang H, Teng L, Fredberg JJ, Guo M (2020) Cell swelling, softening and invasion in a three-dimensional breast cancer model. Nat Phys 16:101–108. 10.1038/s41567-019-0680-8

67. 67. Aureille J, Buffière-Ribot V, Harvey BE, Boyault C, Pernet L, Andersen T, Bacola G, Balland M, Fraboulet S, Van Landeghem L, Guilluy C (2019) Nuclear envelope deformation controls cell cycle progression in response to mechanical force. EMBO Rep 20:1–11. 10.15252/embr.201948084

68. Koushki N, Ghagre A, Srivastava LK, Molter C, Ehrlicher AJ (2023) Nuclear compression regulates YAP spatiotemporal fluctuations in living cells. Proc Natl Acad Sci 120:2017. 10.1073/pnas.2301285120

69. Wakatsuki T, Kolodney MS, Zahalak GI, Elson EL (2000) Cell mechanics studied by a reconstituted model tissue. Biophys J 79:2353–2368. 10.1016/S0006-3495(00)76481-2

70. Steinwachs J, Metzner C, Skodzek K, Lang N, Thievessen I, Mark C, Münster S, Aifantis KE, Fabry B (2016) Three-dimensional force microscopy of cells in biopolymer networks. Nat Methods 13:171–176. 10.1038/nmeth.3685

71. Trappmann B, Gautrot JE, Connelly JT, Strange DGT, Li Y, Oyen ML, Cohen Stuart MA, Boehm H, Li B, Vogel V, Spatz JP, Watt FM, Huck WTS (2012) Extracellular-matrix tethering regulates stem-cell fate. Nat Mater 11:642–649. 10.1038/nmat3339

72. Trepat X, Deng L, An SS, Navajas D, Tschumperlin DJ, Gerthoffer WT, Butler JP, Fredberg JJ (2007) Universal physical responses to stretch in the living cell. Nature 447:592–595. 10.1038/nature05824

73. Krishnan R, Park CY, Lin YC, Mead J, Jaspers RT, Trepat X, Lenormand G, Tambe D, Smolensky A V., Knoll AH, Butler JP, Fredberg JJ (2009) Reinforcement versus fluidization in cytoskeletal mechanoresponsiveness. PLoS One 4:. 10.1371/journal.pone.0005486

74. Fredberg JJ, Inouye D, Miller B, Nathan M, Jafari S, Raboudi SH, Butler JP, Shore SA (1997) Airway smooth muscle, tidal stretches, and dynamically determined contractile states. Am J Respir Crit Care Med 156:1752–1759. 10.1164/ajrccm.156.6.9611016

75. Costa KD, Hucker WJ, Yin FCP (2002) Buckling of actin stress fibers: A new wrinkle in the cytoskeletal tapestry. Cell Motil Cytoskeleton 52:266–274. 10.1002/cm.10056

76. Chaudhuri O, Parekh SH, Fletcher DA (2007) Reversible stress softening of actin networks. Nature 445:295–298. 10.1038/nature05459

77. 77. Vahabi M, Sharma A, Licup AJ, Van Oosten ASG, Galie PA, Janmey PA, Mackintosh FC (2016) Elasticity of fibrous networks under uniaxial prestress. Soft Matter 12:5050–5060. 10.1039/c6sm00606j

78. Murrell MP, Gardel ML (2012) F-actin buckling coordinates contractility and severing in a biomimetic actomyosin cortex. Proc Natl Acad Sci U S A 109:20820–20825. 10.1073/pnas.1214753109

79. McBeath R, Pirone DM, Nelson CM, Bhadriraju K, Chen CS (2004) Cell Shape, Cytoskeletal Tension, and RhoA Regulate Stem Cell Lineage Commitment. Dev Cell 6:483–495. 10.1016/S1534-5807(04)00075-9

80. Chen CS, Mrksich M, Huang S, Whitesides GM, Ingber DE (1997) Geometric Control of Cell Life and Death. Science (80- ) 276:1425–1428. 10.1126/science.276.5317.1425

81. Nelson CM, Khauv D, Bissell MJ, Radisky DC (2008) Change in cell shape is required for matrix metalloproteinase-induced epithelial-mesenchymal transition of mammary epithelial cells. J Cell Biochem 105:25–33. 10.1002/jcb.21821

82. Boudou T, Legant WR, Mu A, Borochin MA, Thavandiran N, Radisic M, Zandstra PW, Epstein JA, Margulies KB, Chen CS (2012) A microfabricated platform to measure and manipulate the mechanics of engineered cardiac microtissues. Tissue Eng - Part A 18:910–919. 10.1089/ten.tea.2011.0341

83. Bonnet-Gonnet C, Belloni L, Cabane B (1994) Osmotic Pressure of Latex Dispersions. Langmuir 10:4012–4021. 10.1021/la00023a019

84. Bouchoux A, Cayemitte PE, Jardin J, Gésan-Guiziou G, Cabane B (2009) Casein micelle dispersions under osmotic stress. Biophys J 96:693–706. 10.1016/j.bpj.2008.10.006

85. 85. Cabane B, Hénon S (2007) Liquides – solutions, dispersions, émulsions, gels, Belin Éduc

86. Thielicke W, Stamhuis EJ (2014) PIVlab – Towards User-friendly, Affordable and Accurate Digital Particle Image Velocimetry in MATLAB. J Open Res Softw 2:. 10.5334/jors.bl

87. Nogueira J, Lecuona A, Rodríguez PA (1997) Data validation, false vectors correction and derived magnitudes calculation on PIV data. Meas Sci Technol 8:1493–1501. 10.1088/0957-0233/8/12/012

88. Westerweel J, Scarano F (2005) Universal outlier detection for PIV data. Exp Fluids 39:1096– 1100. 10.1007/s00348-005-0016-6

89. Garcia D (2010) Robust smoothing of gridded data in one and higher dimensions with missing values. Comput Stat Data Anal 54:1167–1178. 10.1016/j.csda.2009.09.020

90. Tseng Q, Wang I, Duchemin-Pelletier E, Azioune A, Carpi N, Gao J, Filhol O, Piel M, Théry M, Balland M (2011) A new micropatterning method of soft substrates reveals that different tumorigenic signals can promote or reduce cell contraction levels. Lab Chip 11:2231–2240. 10.1039/c0lc00641f

91. Ruppel A, Wörthmüller D, Misiak V, Kelkar M, Wang I, Moreau P, Méry A, Révilloud J, Charras G, Cappello G, Boudou T, Schwarz US, Balland M (2023) Force propagation between epithelial cells depends on active coupling and mechano-structural polarization. Elife 12:1–15. 10.7554/ELIFE.83588

92. Ruppel A, Misiak V, Balland M (2025) An Open-source Python Tool for Traction Force Microscopy on Micropatterned Substrates. Bio-protocol 15:1–15. 10.21769/BioProtoc.5156

93. Sabass B, Gardel ML, Waterman CM, Schwarz US (2008) High resolution traction force microscopy based on experimental and computational advances. Biophys J 94:207–220. 10.1529/biophysj.107.113670

94. 94. Milloud R, Destaing O, de Mets R, Bourrin-Reynard I, Oddou C, Delon A, Wang I, Albigès-Rizo C, Balland M (2017) αvβ3 integrins negatively regulate cellular forces by phosphorylation of its distal NPXY site. Biol Cell 109:127–137. 10.1111/boc.201600041

